# Chloride dynamics alter the input-output properties of neurons

**DOI:** 10.1101/710277

**Authors:** Christopher B. Currin, Andrew J. Trevelyan, Colin J. Akerman, Joseph V. Raimondo

## Abstract

Fast synaptic inhibition is a critical determinant of neuronal output, with subcellular targeting of synaptic inhibition able to exert different transformations of the neuronal input-output function. At the receptor level, synaptic inhibition is primarily mediated by chloride-permeable Type A GABA receptors. Consequently, dynamics in the neuronal chloride concentration can alter the functional properties of inhibitory synapses. How differences in the spatial targeting of inhibitory synapses interact with intracellular chloride dynamics to modulate the input-output function of neurons is not well understood. To address this, we developed computational models of multi-compartment neurons that incorporate experimentally parametrised mechanisms to account for neuronal chloride influx, diffusion, and extrusion. We found that synaptic input (either excitatory, inhibitory, or both) can lead to subcellular variations in chloride concentration, despite a uniform distribution of chloride extrusion mechanisms. Accounting for chloride changes resulted in substantial alterations in the neuronal input-output function. This was particularly the case for peripherally targeted dendritic inhibition where dynamic chloride compromised the ability of inhibition to offset neuronal input-output curves. Our simulations revealed that progressive changes in chloride concentration mean that the neuronal input-output function is not static but varies significantly as a function of the duration of synaptic drive. Finally, we found that the observed effects of dynamic chloride on neuronal output were entirely mediated by changes in the dendritic reversal potential for GABA. Our findings provide a framework for understanding the computational effects of chloride dynamics on dendritically targeted synaptic inhibition.

**Author Summary:** The fundamental unit of computation in the brain is the neuron, whose output reflects information within the brain. A determining factor in the transfer and processing of information in the brain is the modulation of activity by inhibitory synaptic inputs. Fast synaptic inhibition is mediated by the neurotransmitter GABA binding to GABA_A_ receptors, which are permeable to chloride ions. How changes in chloride ion concentration affect neuronal output is an important consideration for information flow in the brain that is currently not being thoroughly investigated. In this research, we used multi-compartmental models of neurons to link the deleterious effects that accumulation of chloride ions can have on inhibitory signalling with changes in neuronal ouput. Together, our results highlight the importance of accounting for changes in chloride concentration in theoretical and computer-based models that seek to explore the computational properties of inhibition.

## Introduction

Neurons in the brain communicate with one another via synaptic signalling, which relies on the activation of receptor proteins that permit rapid transmembrane fluxes of ions. A neuron’s principal operation is to transform synaptic inputs into action potentials. The rate at which action potentials are generated, a neuron’s output firing rate, is a primary means by which neurons encode information (1). The manner in which a neuron integrates rate-coded synaptic input, and transforms it into an output firing rate, is referred to as its input-output function (2).

Synaptic Inhibition is crucial for shaping this input-output transformation (3,4). For example, inhibitory inputs can perform a divisive operation by changing the slope (gain) of the input-output function, or a subtractive operation by offsetting the input-output function to the right (5,6). Previous work has demonstrated that this differential algebraic effect of synaptic inhibition can be realised through targeting inhibitory inputs to different areas of the pyramidal cell. Proximally located inhibition close to the soma can have a divisive effect on the input-output function whilst spatially distributed, dendritically targeted inhibition supports subtractive effects of inhibition (7).

At the receptor level, synaptic inhibition in the brain is primarily mediated by type A γ-aminobutyric acid receptors (GABA_A_Rs), which are selectively permeable to chloride ions (Cl^−^) and, to a lesser extent, bicarbonate ions (HCO_3_^−^) (8). As a result, the reversal potential for GABA_A_Rs (EGABA), is largely a function of the transmembrane Cl^−^ gradient. Together with the membrane potential, this sets the driving force for Cl^−^ flux across these receptors and hence represents a fundamental property of inhibitory signalling (9). Intracellular Cl^−^ concentration ([Cl^−^]_i_), and hence EGABA, can differ between subcellular neuronal compartments, which has been suggested to result from spatial differences in the function or expression of cation-chloride cotransporters in the neuronal membrane (10–12).

In addition to spatial differences in EGABA, inhibitory synapses also exhibit a form of short-term plasticity that involves changes in the ionic driving force for post-synaptic ionotropic receptors driven by short-term alterations in intracellular Cl^−^ concentration (13,14). Ionic plasticity occurs when GABA_A_R-mediated Cl^−^ influx overwhelms local Cl^−^ extrusion via the canonical Cl^−^ extruder KCC2 (15–17).

Previous work has shown how ionic plasticity is regulated on a spatial level with different neuronal subcellular compartments having differing susceptibility to activity-induced chloride accumulation. For example, both experimental and modelling studies have shown how small volume dendritic compartments are particularly prone to ionic plasticity (16,18-21).

Whilst the vast majority of theoretical models of inhibitory signalling and neuronal computation assume static values for EGABA, previous studies have addressed the biophysical underpinnings of Cl^−^ homeostasis and ionic plasticity in neurons (20,22), the effects of neuronal morphologies on Cl^−^ accumulation (19,21,23), and how dynamic Cl^−^ affects neural coding (24). However, how differences in the spatial targeting of synaptic inhibition interact with ionic plasticity to dynamically modulate the input-output function of neurons is not well understood. To address this, we developed computational models of multi-compartment neurons, which incorporated experimentally parametrized mechanisms to account for neuronal Cl^−^ extrusion and ionic plasticity. Firstly, we show that ongoing structured synaptic input (either excitatory, inhibitory, or both) can lead to spatial variations in EGABA independent of differences in Cl^−^ extrusion. Secondly, we find that accounting for dynamic chloride has a significant effect on the ability of dendritically targeted inhibition to offset neuronal input-output curves. Thirdly, we demonstrate that due to ionic plasticity, the neuronal input-output function is not static but varies significantly as a function of time. Finally, we find that the observed effects of dynamic chloride on the neuronal output function are entirely mediated by changes in EGABA. Our results provide a framework for understanding how Cl^−^ dynamics alter the computational properties of spatially targeted synaptic inhibition.

## Results

### Experimental characterisation of neuronal chloride extrusion

To explore the relevance of Cl^−^ dynamics for the computational properties of synaptic inhibition we developed multi-compartment models in NEURON that explicitly accounted for changes in [Cl^−^]_i_ across time and space. As a result, an important parameter within our model which we needed to determine was the dynamics of Cl^−^ extrusion, which is mediated by the canonical Cl^−^ cotransporter in mature neurons, KCC2 (25). We utilised gramicidin perforated patch-clamp recordings in conjunction with optogenetic manipulation of [Cl^−^]_i_ to experimentally characterise Cl^−^ extrusion in hippocampal neurons from mature rat organotypic brain slices (Fig 1A, schematic). Activation of the inward Cl^−^ pump halorhodopsin from *Natronomonas pharaonis* (halorhodopsin, eNpHR3.0 or eNpHR) using a green laser (532 nm) for 15 s resulted in a profound increase in [Cl^−^]_i_, as determined by converted EGABA measurements using gramicidin whole-cell patch-clamp recordings (which do not disturb [Cl^−^]_i_) in conjunction with activation of GABA_A_RS using pressurized ejection of GABA (Fig 1A). [Cl^−^]_i_ recovered to baseline levels over the course of approximately a minute following the 15 s eNpHR photocurrent. This recovery curve could be fitted using a single exponential function and replotted with Cl^−^ extrusion rate (mM/s) as a function of Cl^−^ concentration (Fig 1B). As Cl^−^ extrusion by KCC2 is a function the transmembrane gradient for both K^+^ and Cl^−^, we modelled Cl^−^ extrusion in the manner proposed by Fraser and Huang (26):

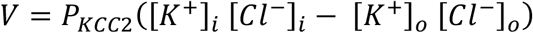

where V is the Cl^−^ extrusion rate and P_KCC2_ is the Cl^−^ extrusion constant, or “pump strength”, of KCC2. The gradient of the line in Fig 1B can be used to calculate P_KCC2_. We measured P_KCC2_ as 1.0 ± 0.16 1/(M⋅s) (mean ± SEM, N = 8). P_KCC2_ was reformulated as a function of surface area (by assuming recorded pyramidal cells had a volume of 1.058 pL with a surface area of 529 × 10^−8^ cm^2^ (27) and current by multiplying by Faraday’s constant, F, such that P_KCC2_ = 1.9297 × 10^−5^ mA/(mM^2^⋅cm^2^). This was then used to calculate the Cl^−^ extrusion rate (V) for each compartment of our model at each point in time depending on the surface area of the compartment concerned and time-varying [Cl^−^]_i_. Combined with mechanisms to describe Cl^−^ influx via both tonic and synaptic activation of GABA_A_RS, as well as Cl^−^ diffusion between compartments (see Materials and Methods), we were able to model how changes in [Cl^−^]_i_ both affects, and is affected by, spatially distributed patterns of synaptic drive.

**Fig 1.**
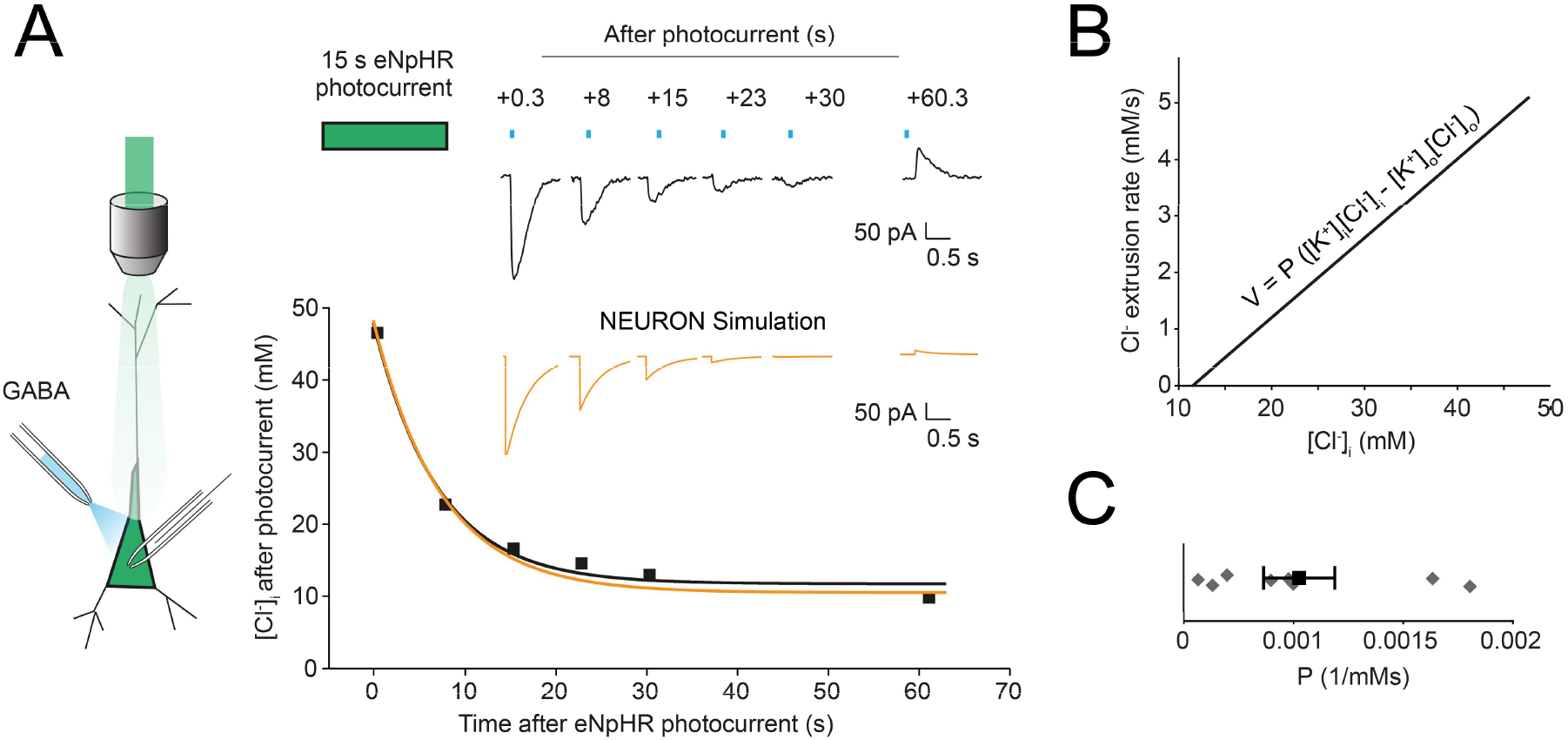
Optogenetic quantification of chloride extrusion in hippocampal neurons. (A) Left, schematic of the experimental setup where a gramicidin perforated patch is made from an eNpHR-expressing hippocampal pyramidal neuron. Green light is delivered via the objective and GABA puffs (blue) directed at the cell soma. Right, top, gramicidin perforated patch voltage-clamp recording from a neuron expressing eNpHR3.0-EYFP. GABA_A_R currents recorded at different times, on different trials following 15s of Cl^−^ load induced by light activation of eNpHR. Right bottom, EGABA and [Cl^−^]_i_ were calculated from each GABA_A_R current (squares, see Materials and methods) and plotted as a function of time after the photocurrent for a single cell. [Cl^−^]_i_ recovery was fitted by a single-exponential function (black). (B) From the data in ‘A’, KCC2 Cl^−^ extrusion rate (V) as a function of [Cl^−^]_i_ was calculated. This allowed for the Cl^−^ extrusion constant, P, to be calculated. (C) Population data from 8 neurons resulted in an average value of P of 0.001 mM^−1^ ⋅ s^−1^. A single compartmental model was then created using the NEURON simulation environment. By accounting for Cl^−^ dynamics including Cl^−^ extrusion via KCC2, a Cl^−^ recovery curve and simulated GABA_A_R currents could be generated (orange curves in ‘A’). These closely match our experimental data.

### Spatial variation in intracellular chloride concentration is affected by the pattern of synaptic drive

In pyramidal neurons, synaptic excitation is predominantly located relatively far from the soma on the branches of the dendrites (28). In contrast, synaptic inhibition either targets proximal structures (soma or proximal dendrites) or is co-located with excitation on peripheral dendrites (28). Extensive evidence exists to suggest that different neuronal subcellular compartments (e.g. soma vs distal dendrites) have different [Cl^−^]_i_ and hence GABAergic inputs to these compartments may have different functional effects. Typically, these differences have been suggested to be driven by differences in the action of cation-chloride cotransporters such as KCC2 (10,11), however, it has also been well described that GABAergic input itself causes differential shifts in [Cl^−^]_i_ depending on the particular subcellular target location (16). What has not been well elucidated is how relative amounts of inhibitory and excitatory synaptic drive may drive spatial variations in [Cl^−^]_i_ despite uniform expression of cation-chloride cotransport.

To investigate this, we generated a multi-compartmental, conductance-based neuron with a short (50 μm), thick (2 μm), ‘proximal’ dendrite connected to a long (500 μm), thin (1.0 μm), ‘distal’ dendrite (see Pouille et al., 2013; Trevelyan & Watkinson, 2005). Synapses were modelled as conductance-based receptors receiving input at a particular input frequency (see Materials and Methods). Excitatory synapses were distally distributed, whilst inhibitory synapses were placed either distally (Fig 2A, inset) or proximally (Fig 2D, inset). Tonic “leak” currents for the three major ions (K^+^:Na^+^:Cl^−^) with relative permeabilities of (1:0.23:0.4) established the steady-state membrane voltage (−71 mV) and resulted in an input resistance of 350 MΩ. In order to select a particular synaptic input structure, the number of excitatory and inhibitory synapses used was determined by tuning the number of synapses, each receiving an input at a mean frequency of 5 Hz from independent Poisson processes, until the neuron produced an output firing rate of 5 Hz (white areas in Fig 2B) under conditions where Cl^−^ was static and held at 4.25 mM. Multiple candidate sets of excitatory-inhibitory (E:I) synapses matched this objective (Fig 2B, blue squares). [Cl^−^]_i_ was then initialized at 4.25 mM but allowed to evolve dynamically over the course of the 1 s simulation as a function of Cl^−^ extrusion by KCC2 (uniformly distributed across all compartments), flux though tonic Cl^−^ receptors, GABAergic synapses, and diffusion between compartments (see Materials and Methods).

**Fig 2.**
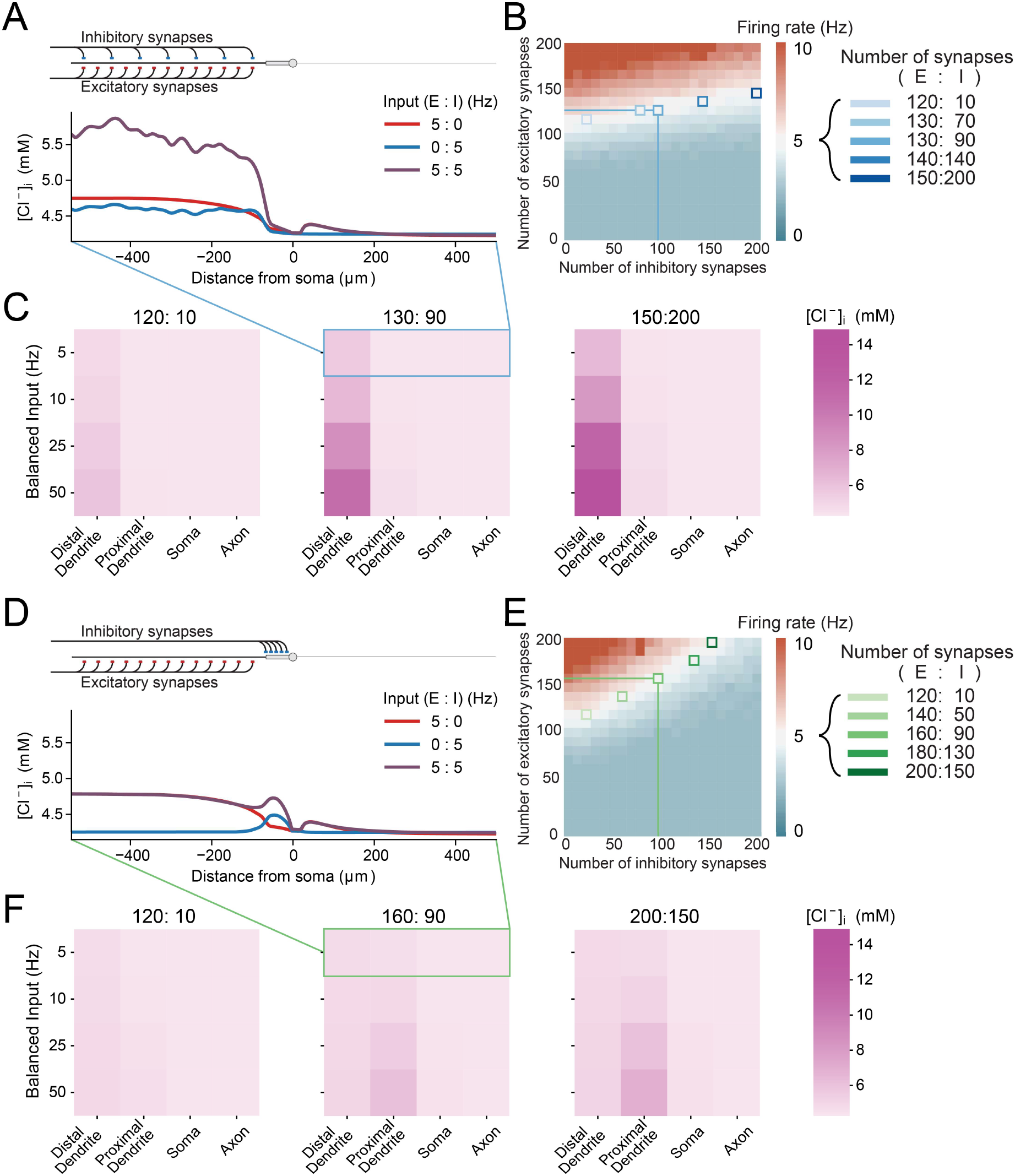
The relative amounts of inhibitory and excitatory drive produce spatial variations in neuronal chloride concentration. (A) Top, schematic depicting the model with an axon, soma, short thick ‘proximal’ dendrite connected to a long thin ‘distal’ dendrite. Both excitatory (red) and inhibitory (blue) synaptic input were evenly distributed across the distal dendrite. Bottom, [Cl^−^]_i_ concentration at the end of the 1 s simulations as a function of distance from the soma. Both inhibitory synaptic input alone (blue trace) and excitatory input alone (red trace) caused selective increases in [Cl^−^]_i_ in the distal dendrite. Balanced input (purple trace) caused an even greater increase in dendritic [Cl^−^]_i_. (B) Heatmap of neuronal firing rates as a function of different numbers of excitatory and inhibitory synapses used to select “balanced” synaptic structures: i.e. pairs of excitatory-inhibitory (E:I) synapse numbers that could transform 5 Hz balanced input into 5 Hz output under conditions of static Cl^−^ (pale squares). (C) Heatmaps of [Cl^−^]_i_ in the distal dendrite, proximal dendrite, soma, and axon at the end of the 1 s simulations when Cl^−^ was allowed to evolve dynamically for three pairs of synaptic numbers reflecting “weak” inhibition (left, E:I - 120:10), “moderate” inhibition (E:I - 130:90) and “strong” inhibition (E:I - 150:200) at different balanced input frequencies. Spatial variations in [Cl^−^]_i_ are dictated by the number of synapses and exaggerated by higher frequencies. (D) Top, schematic as in ‘A’ but with excitatory synapses targeted at the distal dendrite and the inhibitory synapses targeted at the proximal dendrite. Bottom, as in ‘A’, peri-somatic inhibitory synaptic input also produces spatial variations in [Cl^−^]_i_, but of smaller magnitude than when inhibitory synapses are distally located. (E) Heatmap as in ‘B’ but with proximal inhibition. This allowed E:I synapse pairs which produced 5Hz output following 5 Hz input to be identified (pale squares). (F) While functionally similar to synapse pairs with distally targeted inhibition in ‘C’, pairs with proximal inhibition produced more modest changes in [Cl^−^]_i_ as a function of the subcellular domain.

Picking a synaptic structure of 130 excitatory and 90 inhibitory synapses (130:90, E:I), the excitatory synapses alone (Fig 2A, red trace), inhibitory synapses alone (blue trace), or both combined (purple trace) were driven at 5Hz and the final [Cl^−^]_i_ at the end of the 1 s simulation plotted as a function of distance from the soma (Fig 2A). We found that all three modes of synaptic drive produced shifts in [Cl^−^]_i_ in the dendrites relative to other subcellular compartments. Excitatory drive alone was sufficient to cause dendritic shifts in [Cl^−^]_i_ due to an increase in the driving force for Cl^−^ influx via tonic Cl^−^ currents (Fig 2A red trace), which in this case was similar to the [Cl^−^]_i_ shifts driven by inhibitory input alone (blue trace). Interestingly, given balanced input (equal amounts of excitatory and inhibitory input, purple trace), [Cl^−^]_i_ was elevated in the distal compartment more than the sum of the excitatory and inhibitory components. This could be accounted for by the shift in driving force for GABAergic input caused by the EPSPs, even at a low 5 Hz activity.

To further explore how synaptic drive affects the subcellular distribution of [Cl^−^]i, we utilised multiple sets of synaptic structures (Fig 2C) as well as different frequencies of balanced input, including higher frequencies akin to short bursts of feedforward activity. The distal dendrite, receiving all the input, had the highest [Cl^−^]i along the neuron, which was exacerbated by larger balanced input frequencies (50 Hz). This was true for all synapse pairs, but [Cl^−^]i accumulation was most pronounced for 150:200 E:I synapses, given the greater number of GABAergic synapses (Fig 2C, right). These simulation results demonstrate that synaptic drive to the dendrites is an important factor in determining the subcellular distribution of [Cl^−^]_i_ within neurons.

As proximal dendrites tend to have larger diameters, and hence larger volumes we predicted that proximally targeted GABAergic input may have a differing effect on [Cl^−^]_i_. Relative amounts of excitatory and inhibitory synapses were chosen as before to maintain an output of 5Hz following balanced 5 Hz input (Fig 2E, green squares over white). Inhibitory input at 5 Hz caused a small increase in [Cl^−^]_i_ in the proximal dendrite, with minor accumulation in the soma and early part of the distal dendrite due to diffusion (Fig 2D, blue). With excitatory input placed distally and inhibitory input placed proximally, their combined balanced input caused only small accumulations of [Cl^−^]_i_ in the proximal dendrite. [Cl^−^]_i_ accumulation from proximal inhibition was lower overall given the same number of inhibitory synapses and slightly more excitatory synapses (160:90 vs the distal 130:90), despite the same frequencies of synaptic drive (Fig 2F).

Together these results demonstrate that despite a uniform subcellular distribution of Cl^−^ extrusion by KCC2, spatial variations in [Cl^−^]i can arise because of inhomogeneous distribution of synapatic drives across the dendrites. Furthermore, excitatory drive plays an important role in setting the neuron’s [Cl^−^]_i_ by altering the driving force of Cl^−^ flux through both tonic Cl^−^ leak channels and GABA_A_ synapses.

### Dynamic chloride accumulation compromises the effectiveness of inhibition during balanced distal synaptic input

Given the substantial effects of balanced synaptic drive on subcellular [Cl^−^]_i_ we observed in our simplified model of a pyramidal neuron, we next set out to determine how accounting for dynamic Cl^−^ might affect neuronal output given differing frequencies of balanced synaptic drive. As in Fig 2B and E, different sets of excitatory and inhibitory synaptic structures were chosen such that 5 Hz balanced input resulted in a 5 Hz output of the neuron under conditions of static [Cl^−^]_i_. This was done for both proximally targeted inhibitory synapses (green) and distally targeted inhibition (Fig 3).

**Fig 3.**
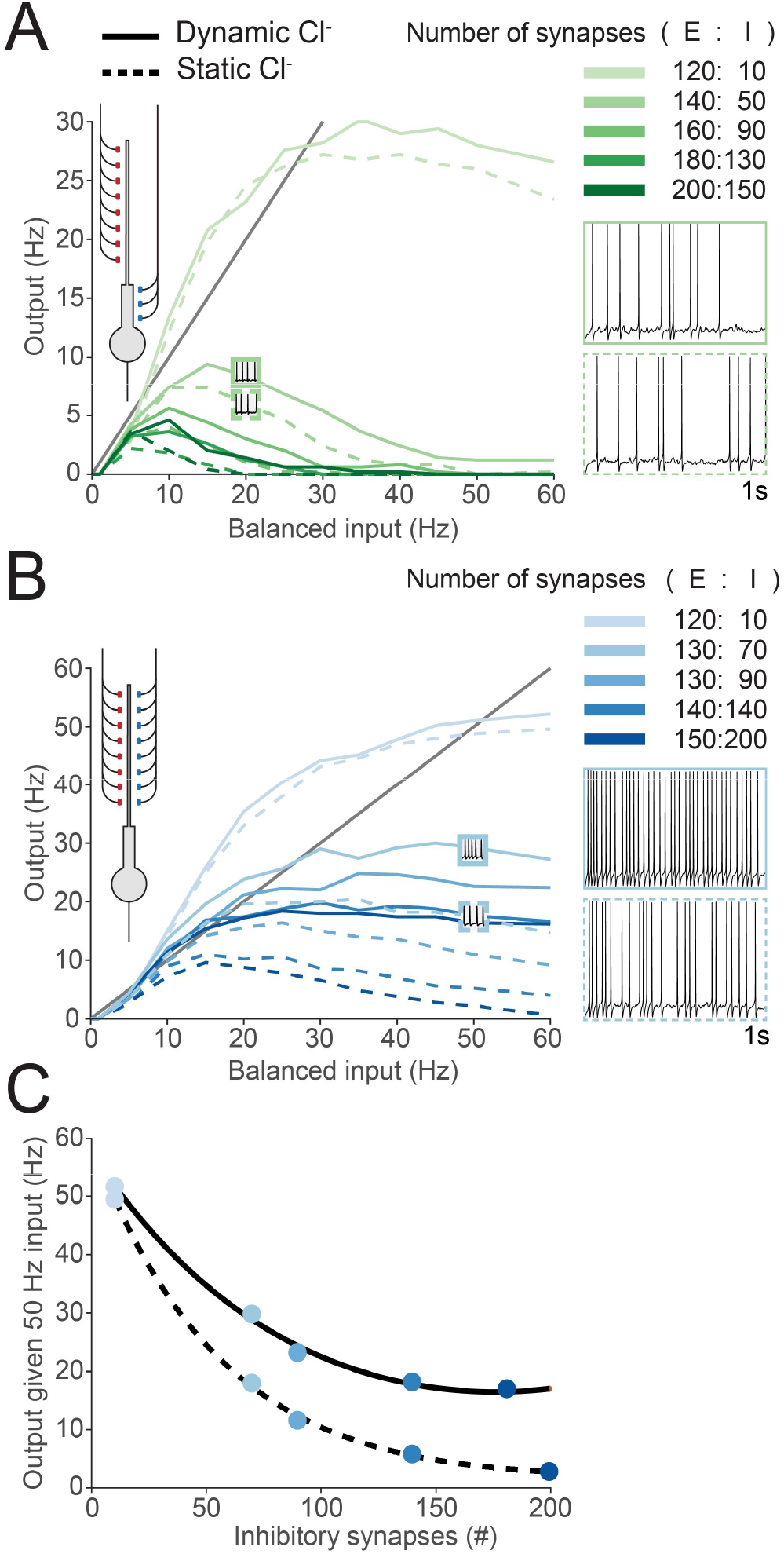
Dynamic chloride accumulation compromises the effectiveness of inhibition during balanced distal synaptic input. (A) Inner left, schematic of the model, depicting excitatory synaptic input targeted toward the distal dendrite and inhibitory synapses targeted at the proximal dendrite. Left, average output firing rates over the course of a 1 s simulation as a function of balanced synaptic input for different pairs of E:I synaptic numbers (shades of green, right). Each pair resulted in a 5 Hz output following 5 Hz balanced input (as in Fig 2). Simulations were performed either with [Cl^−^]_i_ able to vary dynamically ("dynamic chloride", solid lines) or with [Cl^−^]_i_ held at a constant value ("static chloride", dashed lines). Insets with example traces for a simulation run (E:I - 140:50) at 20 Hz input show that accounting for Cl^−^ dynamics results in obvious changes in spike timing. However, chloride dynamics did not result in large changes in output firing rates. (B) Inner left, schematic demonstrating excitatory and inhibitory inputs co-targeted toward the distal dendrite. Left, output firing rates following different balanced input frequencies with different pairs of E:I synaptic numbers (shades of blue, right) as in ‘A’, but with distally targeted inhibition. Dynamic Cl^−^ resulted in large changes to output firing rates (solid vs dashed lines) as well as spike timing (inset, example simulation runs). (C) For neurons receiving high-frequency input (50 Hz), adding more inhibitory synapses above a threshold (≈ 90) did not meaningfully impact the firing rate when Cl^−^ was dynamic. This was not the case for static Cl^−^ where increasing the number of inhibitory synapses continues to decrease output.

For neurons with proximally targeted inhibitory synapses (Fig 3A), the average output firing rate over the course of a 1 s simulation was recorded as a function of balanced synaptic input at multiple frequencies for different synaptic structures (shades of green) and where Cl^−^ was either allowed to evolve dynamically (solid line) or was held static (dashed lines). Immediately apparent was that with a sufficient quantity of proximally targeted inhibitory synapses on neurons receiving higher balanced input frequencies the output firing rate tends to zero. That is, proximal synaptic inhibition curtails neuronal output at higher balanced input frequencies. Although accounting for dynamic Cl^−^ changed the exact timing of spikes compared to the static Cl^−^ condition (Fig 3A, inset), the overall effect of dynamic Cl^−^ on average firing rates was negligible (± 2 Hz difference), likely due to the minimal changes in [Cl^−^]_i_ driven by proximally targeted inhibition as demonstrated in Fig 2E.

In contrast, the results of the previous section suggest that balanced synaptic input with distally targeted inhibitory synapses has significant effects on dendritic [Cl^−^]_i_. To test the functional implications of this, we again measured the output firing rate during the 1 s simulation as a function of multiple balanced input frequencies and different synaptic structures (shades of blue), but now with distally targeted inhibition either with [Cl^−^]_i_ allowed to evolve dynamically, or held static at its initial value (4.25 mM). We observed clear differences in both the timing and rates of action potential generation between simulations run with dynamic vs static Cl^−^ (Fig 3B, inset). Dynamic Cl^−^ resulted in higher output firing rates, particularly for higher balanced input frequencies (Fig 3B). Incorporating Cl^−^ dynamics meant that at high balanced input frequencies, increasing the number of inhibitory synapses beyond a certain number (≈90) had a negligible effect on output firing rates (Fig 3C), with more GABAergic synapses having no additional “inhibitory” effect. That is, when Cl^−^ is dynamic, inhibitory efficacy can not be recovered by simply increasing the number of inhibitory synapses. These data demonstrate that Cl^−^ accumulation compromises the effectiveness of inhibition for controlling output firing rates in response to balanced synaptic input, especially when inhibition is distally targeted.

### Chloride dynamics differentially alter the neuronal input-output function based on inhibitory synaptic location

We next investigated how Cl^−^ dynamics affect the ability of spatially targeted inhibition to alter the input-output function of a neuron. Previous work has demonstrated that dendritically located inhibition, by co-locating inhibition with excitation, can offset a neuron’s input-output curve by increasing the threshold for firing (7,30,31). In this manner, inhibition performs a subtractive operation on the neuronal output function (2). In contrast, inhibition located more proximally than excitation can perform a divisive scaling of the neuronal output, by suppressing action potential generation (2,7,30). How Cl^−^ dynamics affect these operations however is unknown.

To explore this issue, we used the same morphologically simplified model of a pyramidal neuron as before (Fig 4A) to generate a classic input-output curve by plotting average neuronal firing rate during a 1 s simulation as a function of the number of recruited excitatory synapses (Fig 4B). Excitatory synapses were targeted at the distal dendrites and stimulated at 5 Hz from a Poisson distribution. By adding increasing numbers of distally targeted inhibitory synapses (also at 5 Hz) the input-output function could be shifted, or offset, to the right, particularly when [Cl^−^]_i_ was held constant during the simulations (Fig 4B, left). As has been previously described, distal inhibition did not alter the gain or maximum firing rate of the neuron. We observed that allowing Cl^−^ to change during the course of the simulation (dynamic Cl^−^) reduced the ability of distally targeted inhibition to offset the input-output curve (Fig 4B, right).

**Fig 4.**
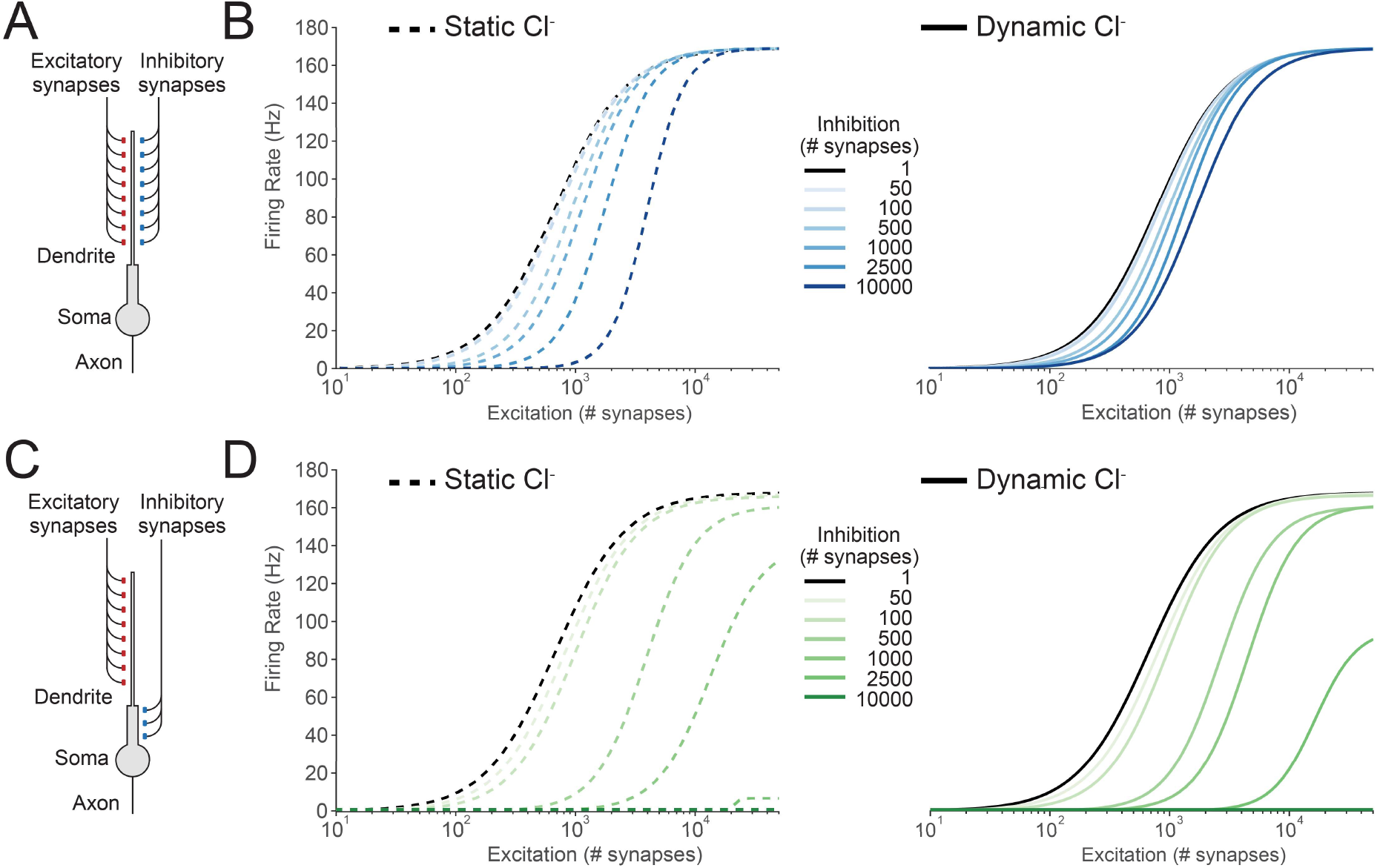
The input-output function of peripherally targeted inhibition is more susceptible to the effects of chloride accumulation than proximally targeted inhibition. (A) Schematic showing the model where both synaptic excitation and inhibition were located on the distal dendrite. Each synapse provided 5 Hz stochastic input to the neuron. (B) Left, average output firing rate over the 1 s simulation as a function of the number of excitatory synapses for different numbers of peripherally located inhibitory synapses, and where Cl^−^ was static (dashed lines). Increasing the number of inhibitory synapses (shades of blue) offset the neuronal input-output curve to the right (a subtractive operation on the input-output function). Right, allowing [Cl^−^]_i_ to vary in the simulations meant that increasing the number of inhibitory synapses had a reduced ability to offset the input-output curve. (C) Schematic, showing a slightly altered version of the model with synaptic inhibition located on the proximal dendrite. (D) Left, as in ‘B’ but inhibition located proximal to excitatory input, i.e. peri-somatically. With static Cl^−^, increasing the number of inhibitory synapses (shades of green) generated divisive gain modulation by both offsetting the threshold and reducing the maximum firing rate of the neuron. Right, output firing rates for the same number of inhibitory synapses were moderately altered with dynamic Cl^−^, yet with sufficient numbers of inhibitory synapses complete suppression of output could still be achieved.

In contrast, simulations with inhibition shifted to a more proximal location (Fig 4C) resulted in inhibition divisively modulating input-output function (Fig 4D). Whilst accounting for dynamic Cl^−^ affected the input-output curves in this condition (e.g. with 2500 inhibitory synapses), the effectiveness of proximal inhibition for divisively modulating the input-output curve remained intact for a broader range of inhibitory synapses than when inhibition was distally located (contrast Fig 4B with Fig 4D). These simulations reveal that Cl^−^ dynamics affect the neuronal input-output function, particularly when inhibition is targeted distally.

### Chloride loading causes rapid degeneration of signalling

A neuron’s input-output function is typically construed as a relatively stable property. However, as demonstrated in Fig 2, continuous synaptic input causes Cl^−^ accumulation and a resultant shift in EGABA, which could progressively affect the input-output function over relatively short timescales. To test this idea, we repeated similar simulations to those described in Fig 4 but with synaptic drive modelled as persistently fluctuating conductances over the full 1 s of each simulation. Instead of measuring the average firing rate over the entire 1 s simulation, we instead calculated the instantaneous firing rate in 20 ms time bins. This allowed us to construct input-output curves as a function of time as the simulations progressed.

We then performed multiple simulations calculating the instantaneous firing rate over time with different levels of excitatory conductances (Fig 5, x-axis of heat maps) and inhibitory conductances (Fig 5, y-axis of heat maps) with [Cl^−^]_i_ as a dynamic variable (Fig 5, upper row) or held static (Fig 5, middle row). By calculating the difference between the dynamic and static Cl^−^ conditions (Fig 5, lower row) we were able to observe how Cl^−^ dynamics affect the input-output curves over time. We noticed that for simulations where neurons received appreciable excitatory and inhibitory synaptic drive, even after as little as 100 ms following the start of the simulation, obvious differences in instantaneous firing rate were evident. That is to say, dendritic inhibition’s control over the neuronal firing rate was progressively compromised over time (Fig 5, lower row). For cases where inhibitory and excitatory input was finely balanced, Cl^−^ dynamics could result in firing rate changes even within 20 ms. In general however, these simulations demonstrate that at very short durations (<20 ms) of synaptic drive, Cl^−^ dynamics do not typically affect the input-output function of neurons, but if this synaptic input continues, changes in [Cl^−^]_i_ progressively affect the neuronal input-output curve, with the potential for large differences in instantaneous firing rate to emerge over the course of one second.

**Fig 5.**
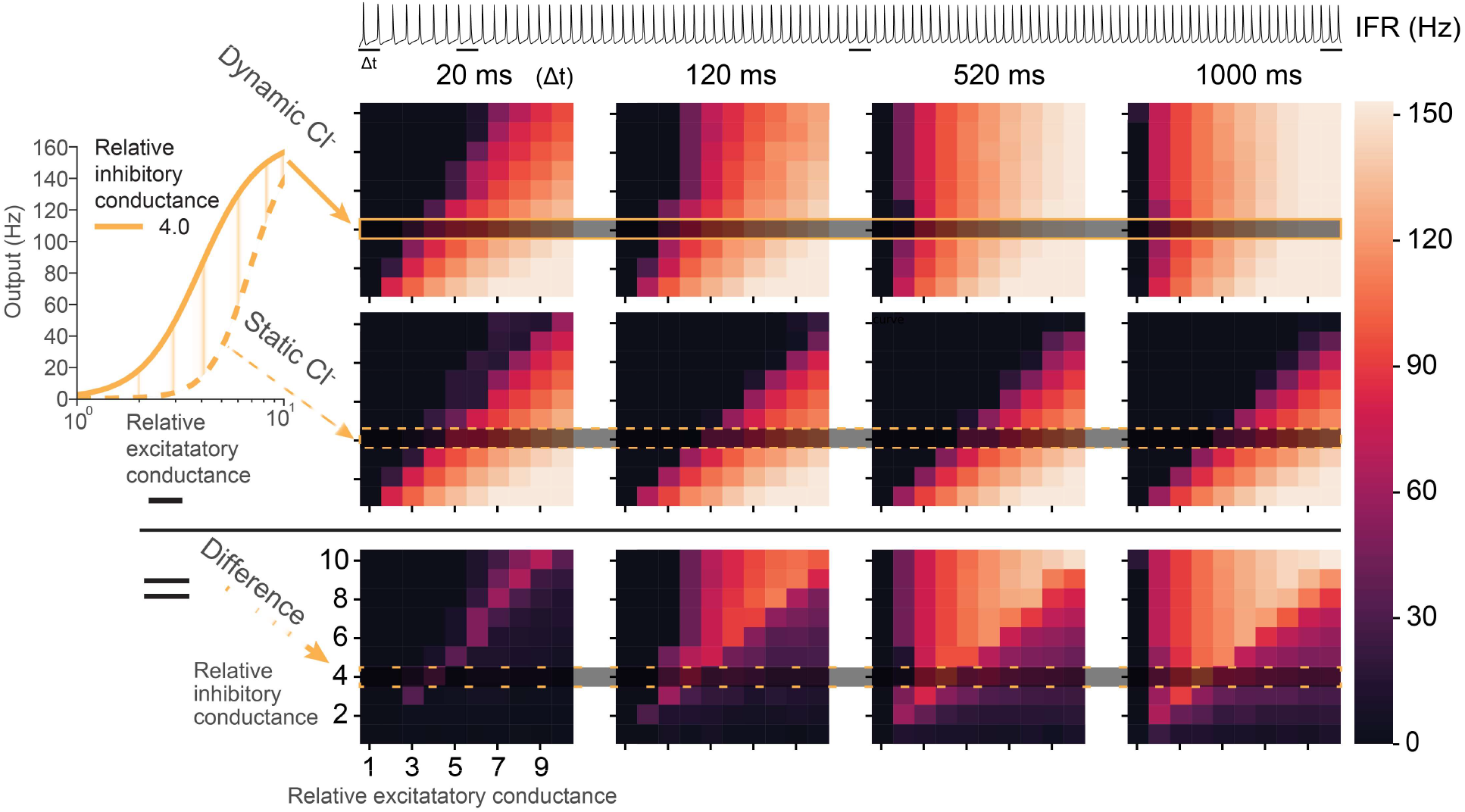
Chloride accumulation has a progressive degenerative effect on the input-output function of neurons. Neuronal input-output curves from instantaneous firing rate (IFR) represented as heatmaps using synapses with continuous fluctuating conductances instead of discrete inputs. Top, an example trace of neuronal membrane potential showing action potential firing averaged over 1 s from a single simulation run. The IFR of the neuron was taken at 20 ms, 120 ms, 520 ms and 1000 ms using a backward integration window (Δt) of 20 ms. Left inset, example input-output curve for simulations with (solid orange trace) and without (dashed orange trace) dynamic Cl^−^. Right, input-output curves represented as heatmaps of IFR for relative excitatory (different columns in an individual heatmap) and inhibitory (different rows in an individual heatmap) conductances. The top row of heatmaps is where simulations were conducted with dynamic Cl^−^. The middle row is for identical simulations with static Cl^−^. The lower row is the difference in IFR between the two. Note how differences in IFR and input-output curves emerge relatively rapidly over the course of 1 second indicating a clear and progressive effect of dynamic Cl^−^ on signalling.

### Quantification of the impact of chloride dynamics in models

In order to determine precisely how Cl^−^ dynamics affect the neuronal input-output function for the case of distally targeted inhibition, we established what we term the “chloride index”. This measure quantifies the extent to which including Cl^−^ dynamics affects the input-output function of a neuron as compared to a condition where [Cl^−^]_i_ is held constant. That is, the “chloride index” is defined as the input-output function shift in a model where Cl^−^ is allowed to evolve dynamically during the simulation (dynamic Cl^−^) as compared to a model with static Cl^−^, relative to the case of no inhibition, as is shown schematically in Fig 6A. This shift is measured from the half-max firing rate, *x*_50_ for a given amount of inhibition *i*.

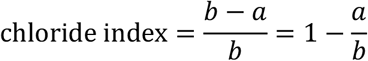

**Fig 6.**
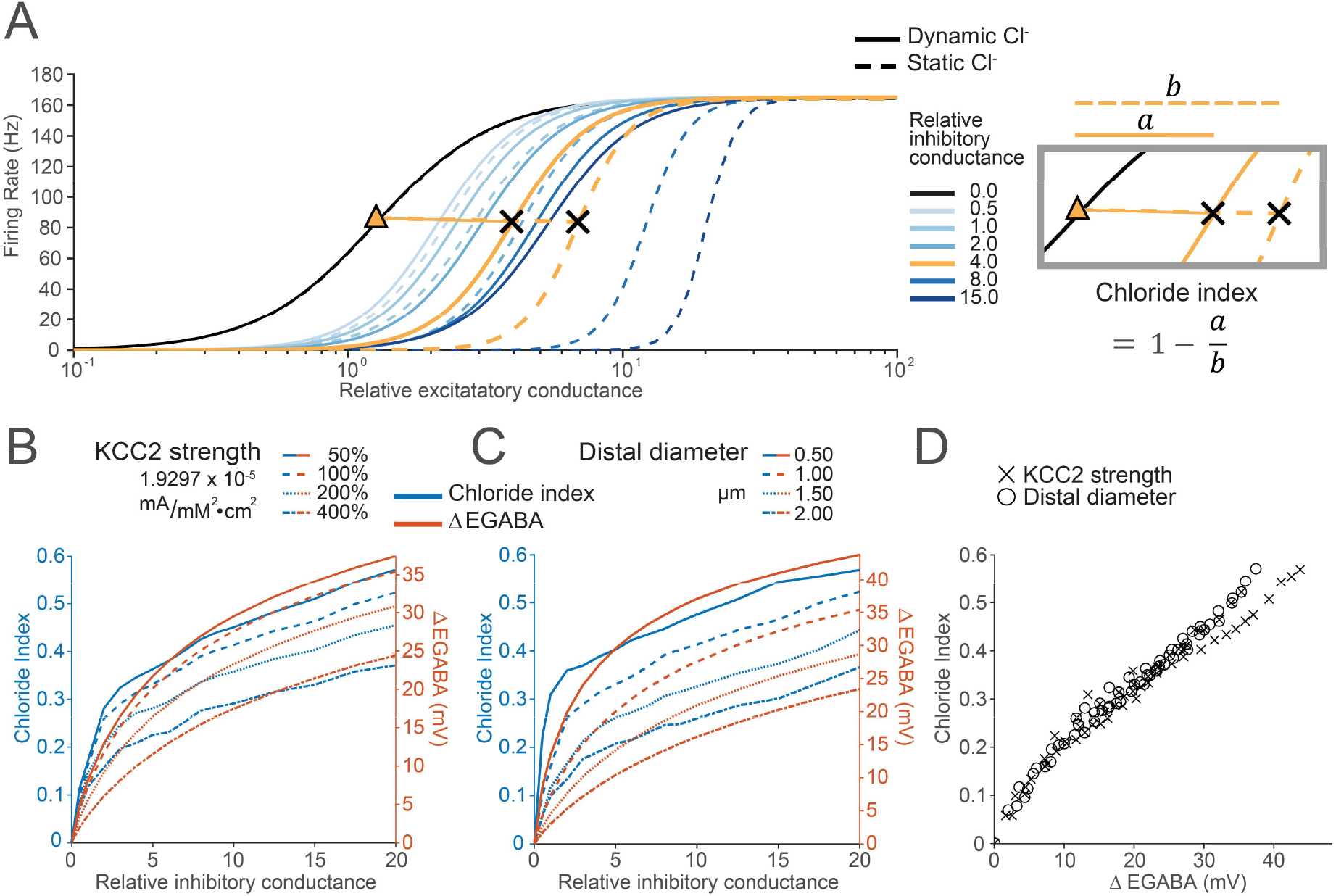
Chloride dynamics shift the neuronal input-output function via changes to EGABA. (A) As in Fig 4 and 5, increasing peripherally targeted inhibition (shades of blue) offset the input-output function, with this effect reduced when Cl^−^ was modelled as dynamic over the course of 1 s simulations (solid lines) as compared to when it was static (dashed lines). The effect of Cl^−^ dynamics on the neuronal input-output curve was quantified using the “chloride index”. Orange traces, and equation (right) demonstrates how the chloride index is calculated using the example of a simulation with a relative inhibitory conductance of 4. Higher Cl^−^ indices (closer to 1) indicate a reduced effect of inhibition due to Cl^−^ being modelled as dynamic instead of static.(B) Chloride index (blue, traces) and the change in dendritic EGABA (∆EGABA, orange traces) after a 1 s simulation for different KCC2 pump strengths (P_KCC2_, 100% = 1.9297 × 10^−5^ mA/mM^2^/cm^2^) as a function of differing levels of peripherally targeted inhibition. Simulations with increased amounts of inhibition resulted in larger shifts in chloride index and ΔEGABA. (C) As in ‘B’ but with P_KCC2_ held at the default level (100%, 1.9297 × 10^−5^ mA/mM^2^/cm^2^) and the diameter of the distal dendrite varied between 0.5 and 2 μm. Small dendritic diameters resulted in larger shifts in chloride index and ΔEGABA during the simulation. (D) Chloride index versus ΔEGABA for all the simulation runs in ‘B’ and ‘C’. The relationship between chloride index - the functional result of accounting for Cl^−^ dynamics - and ΔEGABA is directly proportional and independent of underlying neuronal properties such as KCC2 strength or diameter (r = 0.98464, p < 10^−96^, Pearson correlation, N= 128 simulations).

Where 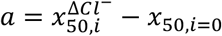

and 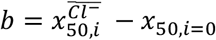.

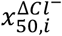 is the half-max firing rate where Cl^−^ is dynamic, 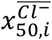 is the half-max firing rate where Cl^−^ is static and *X*_50,i=0_ is the half-max firing rate where there is no inhibition.

In these simulations, synaptic drive was modelled as persistently fluctuating conductances over the full 1 s of each simulation. Firing rates were taken as the average firing rate over the full 1 s simulation. Similarly to the case in Fig 4 and 5, increased inhibitory drive to the distal dendrite offset the input-output curve to the right, with this being reduced by incorporating dynamic Cl^−^. This effect was quantified using the chloride index described above (Fig 6A).

We then repeated these simulations whilst adjusting the model in ways which were predicted to alter the dynamics of Cl^−^ accumulation. First, we systematically adjusted the strength of KCC2 (P_KCC2_), whilst measuring the chloride index as well as EGABA in the distal dendrite (Fig 6B). As demonstrated in Fig 6B, simulations with increased distally targeted inhibition, and in cases where KCC2 activity was reduced, had a greater chloride index because these conditions promote Cl^−^ accumulation. In these simulations, we also measured average EGABA in the distal dendrite at the end of the simulations. We noted a close association between dendritic EGABA (Fig 6B, orange) and the chloride index (Fig 6B, blue). We then repeated these simulations, but instead of adjusting the strength of KCC2, we held KCC2 strength constant and systematically adjusted the diameter of the distal dendrite. Reducing the diameter of the distal dendrite (thereby reducing the volume of this section) also increased the chloride index for any given simulation (Fig 6C). Again, we noticed a close correspondence between the chloride index and EGABA in the distal dendrite. When including all simulations (both adjusting KCC2 or dendritic diameter) we found a very tight, almost linear relationship between the change in EGABA during the simulation in the distal dendrite and the chloride index (r = 0.98464, p < 10^−96^, Pearson correlation, N= 128 simulations, Fig 6D). This clearly demonstrates that regardless of the underlying mechanism, the effect of Cl^−^ dynamics on the input-output function of a neuron can be predicted from the change in EGABA.

## Discussion

The transmembrane Cl^−^ concentration is critically important for regulating the properties of fast synaptic inhibition in the brain. Here we have used computational models of morphologically simplified, multi-compartment neurons that incorporated experimentally parametrised neuronal Cl^−^ extrusion in order to provide a framework for investigating the impact of Cl^−^ dynamics on the input-output properties of neurons. We found that continuous excitatory or inhibitory synaptic input can result in subcellular spatial differences in [Cl^−^]_i_, and that shifts in [Cl^−^]_i_ during synaptic input can affect the input-output properties of neurons.

The idea that [Cl^−^]_i_ can differ spatially within a particular neuron, depending on the subcellular compartment concerned, is important for understanding the effect of spatially targeted synaptic inhibition (10), as well as the consequences of the latest optogenetic silencing strategies which utilise light-activated Cl^−^ channels (32). Many interacting variables determine resting [Cl^−^]_i_ in neurons (20,22), but subcellular differences [Cl^−^]_i_ are typically attributed to differences in the activity of cation-chloride cotransporters such as KCC2 or NKCC1. In support of previous work (20,21), we show that the relative amounts of inhibitory and excitatory drive can also produce spatial variations in [Cl^−^]_i_ despite uniform Cl^−^ extrusion capacity. Notably, we show that even dendritic excitatory synaptic input alone is sufficient to drive shifts [Cl^−^]_i_ by changing the driving force for Cl^−^ flux via tonic Cl^−^ channels. This framework will be useful for experimentalists interpreting data collected using the latest tools to observe subcellular [Cl^−^]_i_ *in vivo* (33), where different activity states will likely affect synaptic drive onto neurons.

Previous experimental and computational work has demonstrated that for a given Cl^−^ load via activated synaptic GABA_A_Rs, Cl^−^ accumulation occurs more readily on smaller volume peripheral dendrites than more proximal dendritic and somatic compartments (16,18,19,21). The effect of this on the input-output properties of neurons, however, has not previously been explored. We found that accounting for Cl^−^ dynamics has a more pronounced effect on dendritically targeted inhibition than proximally located inhibition. This is for several intuitive reasons. Firstly, inhibitory inputs occur on smaller volume structures, which means that for a given Cl^−^ load, there is a greater shift in [Cl^−^]_i_ and EGABA. Secondly, inhibitory inputs are collocated with excitatory inputs so the driving force for Cl^−^ influx is typically greater for the combined excitatory and inhibitory synaptic drive. The result is that the effectiveness of distal inhibition is more susceptible to the weakening effect of Cl^−^ accumulation than proximal inhibition and may be part of the reason why the bulk of hippocampal and cortical pyramidal neuron inhibitory synapses are typically located on their proximal dendrites and somata (28).

In our models, including Cl^−^ dynamics reduced the ability of distally targeted inhibition to offset the neuronal input-output function. That is, it reduced the ability of distally targeted inhibition to perform a subtractive operation. And because greater inhibitory synaptic drive enhances Cl^−^ accumulation and counterintuitively weakens inhibition further, it is not possible to overcome this inhibitory weakening by increasing the strength of distal inhibitory synaptic drive. That is, beyond a certain level, simply increasing GABAergic synaptic activation had no additional “inhibitory” effect. We also show that this perturbation of the input-output function happens progressively over time. Particularly under conditions where inhibitory and excitatory input is finely balanced, Cl^−^ dynamics can rapidly and substantially alter instantaneous firing rates. This underscores the fact that even for neurons with a fixed structure of synaptic input, dynamic Cl^−^ progressively alters the neuronal input-output function. This supports previous work, using single-compartment neurons, that demonstrates how, and under which paradigms, Cl^−^ dynamics can degrade neuronal coding (24). By using a measure we termed the “chloride index” we were able to demonstrate that across various conditions (i.e. different KCC2 pump strengths or dendritic diameters) the shift in dendritic EGABA during synaptic input predicts how Cl^−^ dynamics affect the neuronal input-output function.

Whilst we have demonstrated that Cl^−^ dynamics compromise the ability of distally targeted inhibition to control neuronal output in the form of action potential generation at the soma and axon, in some cell types, particularly pyramidal neurons, it is clear that dendrites host active conductances and that non-linear input summation and integration also occurs within the dendritic tree itself (34,35). Given this, it is likely that an important function of peripherally targeted inhibition is to control local non¬linear summation of excitatory input, which may be more resistant to the effects of Cl^−^ dynamics. Future work will need to determine how Cl^−^ dynamics affects, and is affected by, active dendritic conductances.

We have shown how changes in [Cl^−^]_i_ have significant importance for the control of neuronal output. This is predicted to have important implications for the regulation of network excitability under both physiological and pathological conditions (36). As an extreme example, the intense, simultaneous excitatory and inhibitory input received by pyramidal neurons during epileptic seizures results in severe Cl^−^ accumulation that initially weakens inhibition before resulting in paradoxical GABAergic excitation that perpetuates epileptiform activity (37,38). However, as we demonstrate here, for the case of distally targeted dendritic inhibition, even small activity-driven changes in dendritic [Cl^−^]_i_ can affect neuronal output, with likely implications for network activity. Together, our results highlight the importance of accounting for changes in Cl^−^ concentration in theoretical and computer-based models that seek to explore the computational properties of dendritic inhibition.

## Methods

### Experimental procedures

Rat organotypic hippocampal slice cultures were prepared using a method similar to that described by Stoppini and colleagues (39). 7 day old male Wistar rats were killed in accordance with the UK Animals Scientific Procedures Act 1986. The brains were extracted and placed in cold (4 °C) Geys Balanced Salt Solution (GBSS), supplemented with D-glucose (34.7 mM). The hemispheres were separated and individual hippocampi were removed and immediately sectioned into 350 μm thick slices on a McIlwain tissue chopper. Slices were rinsed in cold dissection media, placed onto Millicell-CM membranes and maintained in culture media containing 25 % EBSS, 50 % MEM, 25 % heat-inactivated horse serum, glucose, and B27 (Invitrogen). Slices were incubated at 36 °C in a 5 % CO_2_-humidified incubator before transfection. Neurons were biolistically transfected after 5-6 days *in vitro* using a Helios Gene Gun (120 psi; Bio-Rad) with pLenti-hSyn-eNpHR3.0-EYFP (eNpHR3.0 fused to EYFP and driven by the human synapsin I promoter, generously provided by the Deisseroth lab, Stanford University). 50 μg of target DNA was precipitated onto 25 mg of 1.6 μm diameter gold microcarriers and bullets generated in accordance with the manufacturer’s instructions (Bio-Rad). At the time of recording, transfected neurons were equivalent to postnatal day 14-17. Previous work has shown that the pyramidal neurons in the organotypic hippocampal brain slice have mature and stable Cl^−^ homeostasis mechanisms, as evidenced by their hyperpolarizing EGABA (40,41) and that KCC2 is the major active Cl^−^ transporter in these neurons as EGABA is affected by KCC2-blocking drugs, but not by NKCC1-blocking drugs (42).

Hippocampal slices were transferred to the recording chamber and continuously superfused with 95 % O_2_ - 5 % CO_2_ oxygenated artificial cerebrospinal fluid (ACSF), heated to 30 °C. The ACSF was composed of (in mM) NaCl (120), KCl (3), MgCl_2_ (2), CaCl_2_ (2), NaH_2_PO_4_ (1.2); NaHCO_3_ (23); D-Glucose (11) and the pH was adjusted to be between 7.38 and 7.42 using NaOH (0.1 mM). Glutamatergic receptors and GABA_B_RS were blocked with kynurenic acid (2 mM) and CGP55845 (5 μM). Neurons within the pyramidal cell layer of the CA1 and CA3 regions of the hippocampus were visualized under a 20x water-immersion objective (Olympus) and targeted for recording. Patch pipettes of 3-7 ΜΩ tip resistance were pulled from filamental borosilicate glass capillaries with an outer diameter of 1.2 mm and an inner diameter of 0.69 mm (Harvard Apparatus Ltd, Hertfordshire, UK), using a horizontal puller (Sutter). For gramicidin perforated patch-clamp recordings(43), pipettes were filled with a high KCl internal solution whose composition was (in mM): KCl (135), Na_2_ATP (4), 0.3 Na_3_GTP (0.3), MgCl_2_ (2), and HEPES (10). Gramicidin (Calbiochem) was dissolved in dimethylsulfoxide to achieve a stock solution of 4 mg/ml. This was then diluted in internal solution immediately prior to experimentation (10 min before attaining a patch) to achieve a final concentration of 80 μg/ml. The resulting solution was vortexed for 1 min, sonicated for 30 s and then filtered with a 0.45 μm pore cellulose acetate membrane filter (Nalgene). The osmolarity of internal solutions was adjusted to 290 mOsM and the pH was adjusted to 7.38 with KOH.

GABA_A_RS were activated by delivering short ‘puffs’ of GABA (100 μM) in the presence of glutamate receptor blockers and GABA_B_R blockers (see above). The agonist was applied via a patch pipette positioned close to the soma and connected to a picospritzer (20 psi for 20ms; General Valve). To measure [Cl^−^]_i_ recovery following a photocurrent-induced [Cl^−^]_i_ load, it was important to estimate EGABA, and hence [Cl^−^]_i_, from single GABA_A_R currents. To achieve this, the resting EGABA and GABA_A_R conductance (gGABA) were calculated before each experiment and these values were then used to estimate EGABA for a single GABA_A_R current by assuming a consistent gGABA across GABA puffs and solving the equation 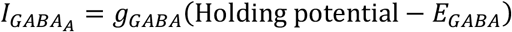. EGABA was converted to [Cl^−^]_i_ using the Nernst equation and assuming a GABA_A_R Cl^−^ to HCO_3_^−^ permeability ratio of 4 to 1, with [HCO_3_^−^]_i_ = 10 mM and [HCO_3_^−^]_o_ = 25 mM. Photoactivation of NpHR to generate 15 s Cl^−^ photocurrents was achieved via a diode-pumped solid state (DPSS) laser, with a maximum output of 35 mW and a peak at 532 nm (Shanghai Laser Optic Century). The laser was attenuated via a 5 % neutral density and coupled to a 1000 μm diameter mulitimode optic fiber via a collimating lens (Thorlabs).

### Modelling

A multi-compartmental, conductance-based neuron model was constructed in NEURON (44). The neuron was constructed as a soma extending to a short (50 μm), thick (2 μm), ‘proximal’ dendrite connected to a long (500 μm), thin (1.0 μm), ‘distal’ dendrite as in (7,29).

Spikes were recorded from the end of the 500 μm long axon. The neuronal morphology used is depicted to scale in Fig 1. The membrane potential was updated at each time step according to transmembrane ion currents from channels and pumps, taking into account the membrane capacitance, together calculated as 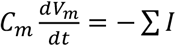. Some simulations used NEURON’s variable time step integrator, ‘CVODE’, but the default time step of 0.025 ms was typically used.

Synapses were modelled as either conductance-based receptors receiving input from independent stochastic Poisson processes (ξ = 1) providing an input frequency (’f-in’) or as persistent fluctuating conductances (’gclamp’). The reversal potentials of excitatory and inhibitory synapses matched those in Pouille et al. (2013): E_excitation_ = 0 mV; E_inhibition_ = −74 mV. Excitatory f-in synapses were modelled as NMDA receptors (τ_decay_ = 100 ms), with Mg^2+^ dependence on membrane voltage (45). F-in synapses were modelled based on the simple conceptual model of transmitter-receptor interaction where transmitter binds to a closed receptor channel to produce an open channel with a forward binding rate α and backward unbinding rate β (46)(see Table 1). The g_max_ values for NMDA and GABA_A_ receptors were 100 pS and 350 pS, respectively, and strength increased by adding synapses. Gclamp synapses had their strength increased by multiplying their baseline passive conductance, 0.0001 μS, by a factor (‘relative conductance’). Fluctuations in conductance, ‘noise’, was set as a coefficient of variance of 0.1. Excitatory synapses were placed on the distal dendrite. Inhibitory synapses were placed either on the proximal or distal dendrite. The number of synapses varied by experiment but were evenly distributed along the respective dendrite.

**Table 1:**
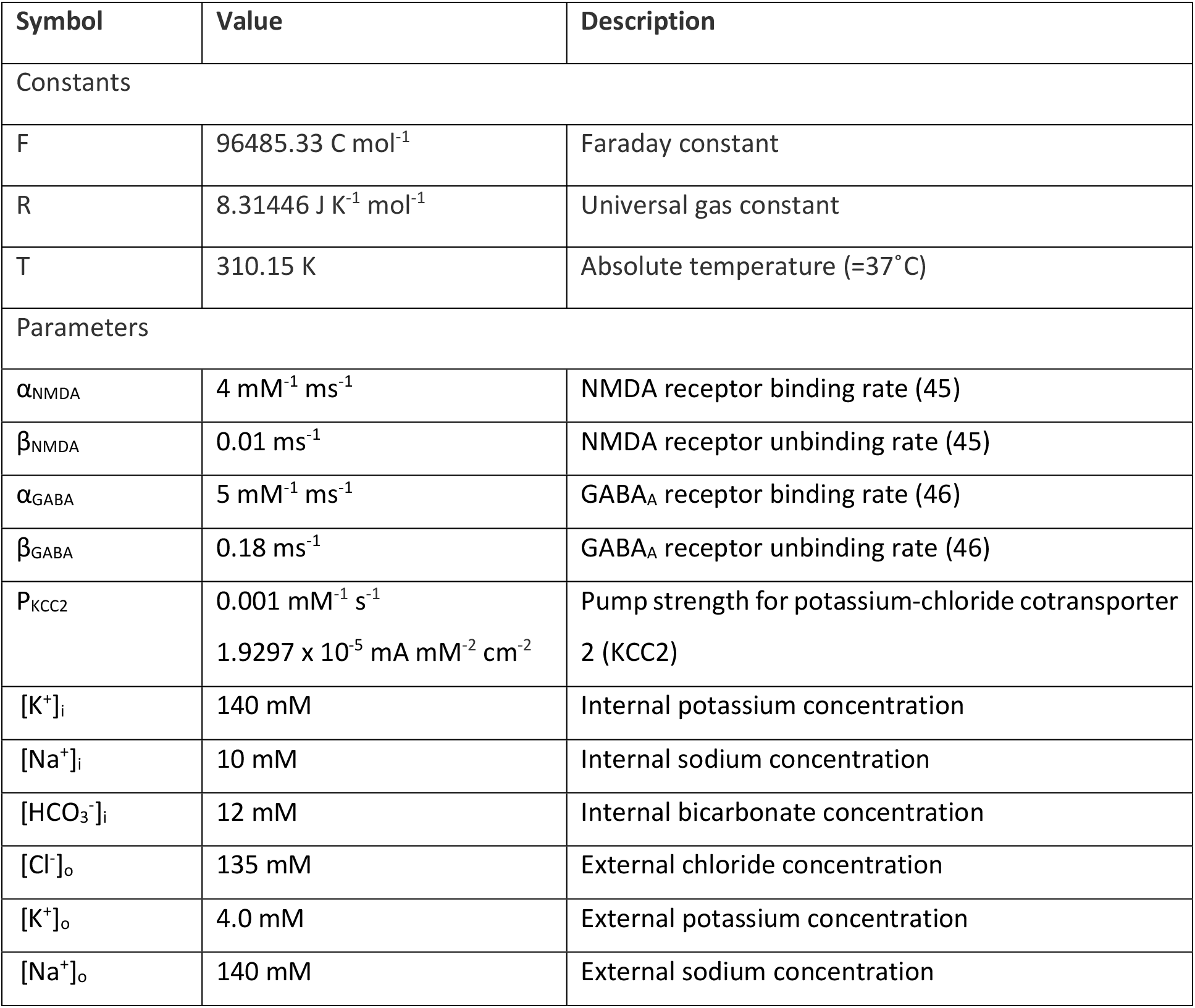
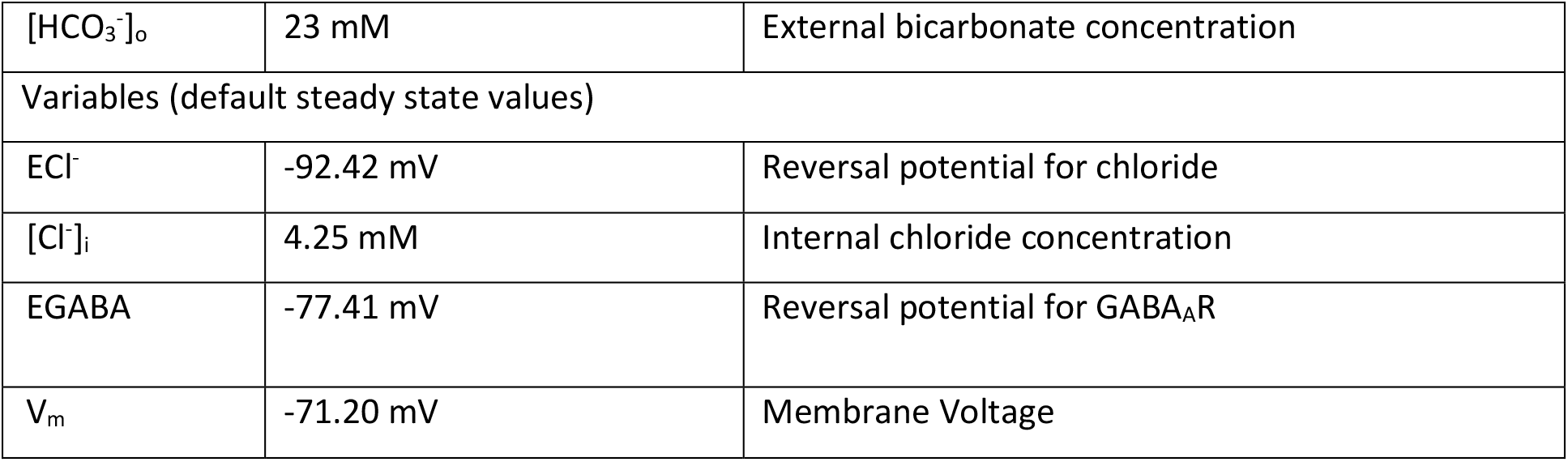
Constants, parameters, and default steady state values for variables

**Table 2:** Abbreviations

|  |  |
| --- | --- |
| Cl<sup>-</sup> | Chloride ion(s) |
| GABA<sub>A</sub>R | Type A γ-aminobutyric acid receptor |
| HCO<sub>3</sub><sup>-</sup> | Bicarbonate ion(s) |

GABA_A_ receptors, regardless of f-in or gclamp, were modelled as being selectively permeable to both Cl^−^ and HCO_3_^−^ ions (4:1 ratio) and used the Ohmic formulation for current 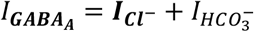, where 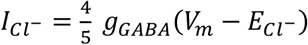 and 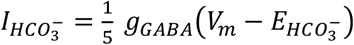. Where applicable, the reversal potential for chloride was updated throughout the simulation using the Nernst equation, 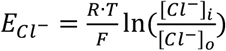. The reversal potential for HCO_3_^−^ was held constant, and EGABA calculated as 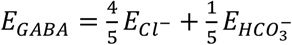.

The soma and axon had channels with Hodgkin-Huxley style kinetics, as in (29), governing the sodium (Na^+^) channel, 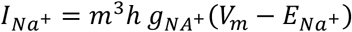; potassium (K^+^) channel, 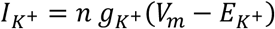; potassium M-current responsible for the adaptation of firing rate and the afterhyperpolarization, 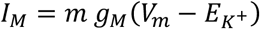; and leak conductances fitted for an input resistance of 350 ΜΩ with a K^+^:Na^+^:Cl^−^ ratio of 1:0.23:0.4. Leak conductances were also in the dendrites and, with KCC2, set the resting [Cl^−^]_i_ to 4.25 mM. Unless stated as ‘static chloride’, [Cl^−^]_i_ was allowed to vary (‘dynamic Cl^−^’). Transmembrane Cl^−^ fluxes due to Cl^−^ currents through GABA_A_RS, KCC2 co-transporter (as in Fig 1), as well as changes due to longitudinal diffusion, were calculated as

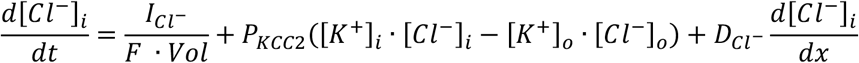

where P_KCC2_ is the “pump strength” of chloride extrusion, **D_cl_-** is the diffusion coefficient for chloride in water, F is Faraday’s constant, Vol is the volume of the compartment, and x is the longitudinal distance between the midpoint of compartments (see Table 1 for parameter values).

The firing rate of a neuron was determined by counting the number of action potentials at the tip of the axon over a 1000 millisecond period. Where applicable, the instantaneous firing rate (IFR) was calculated as

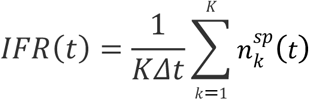

where ***K*** (number of trials) = 5, *∆t* (time bin) = 20 ms, and 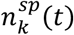 was the number of spikes recorded in trial **k** over the time window [*t* −∆*t*, *t*].

## Supporting Information

### S1. Converting KCC2 “pump strength” parameter from 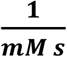 to 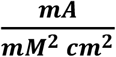

For a cylinder, volume = *L* π *r*^2^ and *surface area = 2L π r*, where *L* and *r* are the length and radius, respectively. Thus, the extrusion constant can be converted as such:

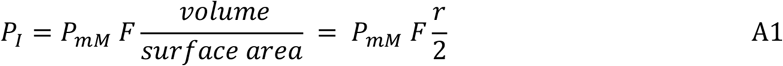

where *P*_*I*_ is the extrusion constant in 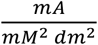, *P*_*mM*_ is the extrusion constant in 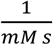, *F* is the Faraday constant in 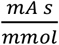.

Checking units, it is convenient to keep the units for *volume* and *surface area*,

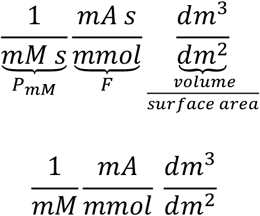

Remembering that 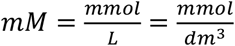,

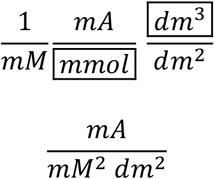

For units in *cm*^2^, as typically used in current per area, 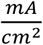,

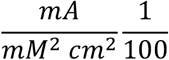

For 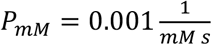 and a neuron segment with *r =* 4 *um*, 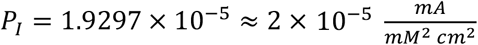.

